# *De novo* purine nucleotide biosynthesis mediated by amidophosphoribosyl transferase is required for conidiation and essential for the successful host colonization of *Magnaporthe oryzae*

**DOI:** 10.1101/2021.02.28.433215

**Authors:** Osakina Aron, Min Wang, Jiayuan Guo, Jagero Frankline Otieno, Qussai Zuriegat, Songmao Lu, Zonghua Wang, Wei Tang

**Affiliations:** Fujian University Key Laboratory for Plant-Microbe Interaction, College of Life Science, Fujian Agriculture and Forestry University, Fuzhou, 350002, China; State Key Laboratory of Ecological Pest Control for Fujian and Taiwan Crops, College of Plant Protection, Fujian Agriculture and Forestry University, Fuzhou 350002, China; Marine and Agricultural Biotechnology Laboratory, Institute of Oceanography, Minjiang University, Fuzhou 350108, China

**Keywords:** *Magnaporthe oryzae*, amidophosphoribosyltransferase, purine nucleotide biosynthesis, pathogenicity defect

## Abstract

Amidophosphoribosyl transferase catalyzes the first step of the purine nucleotide biosynthesis by converting 5-phosphoribosyl-1-pyrophosphate into 5-phosphoribosyl-1-amine. In this study, we identified and characterized the functions of MoAde4, an ortholog of yeast Ade4 in the rice blast fungus. MoAde4 is a 537-amino acid protein containing the GATase_6 and pribosyltran domains. Quantitative real-time PCR analysis showed *MoADE4* transcripts were highly expressed during conidiation, early-infection, and late-infection stages of the fungus. Disruption of *MoADE4* gene resulted in Δ*Moade4* mutant exhibiting adenine, adenosine, and hypoxanthine auxotrophy on MM. Conidia quantification assays showed Δ*Moade4* mutant was significantly reduced in sporulation. The conidia of Δ*Moade4* mutant could still form appressoria but mostly failed to penetrate the rice cuticle. Pathogenicity test showed Δ*Moade4* was completely nonpathogenic on rice and barley leaves which was attributed by failure of its infectious hyphae to colonize the host cells. The Δ*Moade4* was defective in induction of strong host immunity and had its purine transporter genes repressed during in planta infection. Addition of exogenous adenine partially rescued conidiation and pathogenicity defects of the Δ*Moade4* mutant on the barley and rice leaves. Localization assays showed that MoAde4 is located in the cytoplasm. Taken together, our results demonstrate that purine biosynthesis orchestrated by MoAde4 is required for fungal development, conidiation, more importantly, we found it to be essential for fungal pathogenicity not because of the appressorial formation, but appressorium penetration and host colonization during the plant infection of *M. oryzae*. Thus this findings suggests that purine biosynthesis could act as an important target for combating recalcitrant plant fungal pathogens.

## INTRODUCTION

Rice blast, caused by *Magnaporthe oryzae*, is one of the most serious rice diseases worldwide (H. Zhang, Zheng, & Zhang, 2016). Global rice production is under constant threat from rice blast disease which could result in an annual loss of rice yields for consumption for more than half of the global population (Talbot, 2003; Zeigler, 1998). Disease initiation occurs when *M. oryzae* conidia land on rice leaves and differentiate into a highly specialized dome-shaped infectious structure called appressorium (Osés-Ruiz, Sakulkoo, Littlejohn, Martin-Urdiroz, & Talbot, 2017). Upon maturation, enormous turgor pressure is built up within the appressorium, which is important in forcing the penetration peg to physically puncture the plant cuticle (de Jong, McCormack, Smirnoff, & Talbot, 1997). While inside the host cells, infectious hyphae (IH) differentiate and rapidly colonize the plant cells, and this proceeds within a new disease cycle occurring in 5–7 days (Foster, Ryder, Kershaw, & Talbot, 2017).

Rice blast fungus has developed some regulatory system that enables the infection cycle to be completed. These regulatory systems comprises of the carbon catabolite repression (CCR) and nitrogen metabolite repression (NMR) that respectively ensures utilization of carbon (glucose) and preferred sources of nitrogen (ammonium and L-glutamine) (Fernandez et al., 2012; Wilson et al., 2012; Wilson et al., 2007; Wilson & Talbot, 2009). Many enzymes involved in metabolisms such as MoIlv2 and MoIlv6, MoLeu1, MoStr3, and MoMet13, were characterized and found to be involved in the development and infection of rice blast fungus (Du et al., 2013; Tang et al., 2019; Wilson et al., 2012; Yan et al., 2013). Clearly understanding the molecular mechanisms involved in the development and pathogenicity of *M. oryzae* will assist in research geared towards antifungal development (Wilson & Talbot, 2009). Purine nucleotides are essential metabolites for cellular physiology as they constitute structural components of nucleic acids DNA and RNA, energy carriers (i.e. ATP and GTP), and enzyme cofactors (i.e. NAD+ and NADP+). Furthermore, purine nucleotides are important for the biosynthesis of several amino acids and vitamins such as folic acid and riboflavin, which are of significant value in the biotechnology industry (Abbas & Sibirny, 2011; Peifer et al., 2012). These molecules are therefore crucial for all known forms of life. The *de novo* purine biosynthesis pathway occurs in a 10-step enzymatic process and involves the conversion of PRPP (5-phosphoribosyl-α-1-pyrophosphate) into the first purine nucleotide (IMP) (Liechti & Goldberg, 2012). IMP serves as a branching point in the biosynthetic pathway and is transformed into AMP or GMP in two additional steps (Ljungdahl & Daignan-Fornier, 2012). An alternative salvage pathway exits, which generates GMP from guanine and AMP from adenine directly, or indirectly generate IMP or XMP from hypoxanthine and xanthine. These nucleotides are then incorporated into the nucleotide biosynthesis pathway for the generation of GMP or AMP (Kappock, Ealick, & Stubbe, 2000).

Previous studies have demonstrated the importance of purine nucleotide in various organisms. In *Drosophila*, the biosynthesis of purine nucleotide synthesis is required for metamorphosis (Ji & Clark, 2006). In humans, disorders resulting from dysfunctional purine nucleotide biosynthesis have been reported (Balasubramaniam, Duley, & Christodoulou, 2014; Jinnah & Van Den Berghe, 2013; Nyhan, 2005). In microorganisms as *Escherichia coli* and *Saccharomyces cerevisiae*, the impaired biosynthesis of purine nucleotides leads to auxotrophy (Beckwith, 1996; Kokina, Kibilds, & Liepins, 2014). Although purine nucleotide biosynthesis was shown to be involved in the development and infection of rice blast fungus (Fernandez, Yang, Cornwell, Wright, & Wilson, 2013). The exact role of MoAde4, the enzyme that catalyzes the first rate-limiting reaction in *de novo* biosynthetic pathway remains unclear. Here, we identified and characterized the MoAde4 in rice blast fungus *M. oryzae*. We have confirmed that *de novo* purine nucleotide biosynthesis mediated by MoAde4 is crucial for conidiation, and pathogenicity of rice blast fungus.

## Results

### Identification of MoAde4 in *M. oryzae*

To identify MoAde4, the Ade4 amino acid sequence of *Saccharomyces cerevisiae* was used as query for BlastP search in the Kyoto Encyclopedia of Genes and Genome (KEGG) database of *M. oryzae* (http://www.kegg.jp/kegg-bin/show_organism?org=mgr). The obtained MoAde4 amino acid sequence was then used for a Pfam-based domain prediction. Domain analysis revealed that MoAde4 contains GATase_6 and pribosyltran domains (Fig. 1A). Additional domain prediction from the other fungi, including *Ustilago maydis* (Um), *Saccharomyces cerevisiae* (Sc), *Fusarium graminearum* (Fg), *Sclerotinia sclerotiorum* (Ss), *Neurospora crassa* (Nc), *Fusarium oxysporum* (Fo), *Trichoderma reesei* (Tr), *Aspergillus flavus* (Af) and *Aspergillus niger* revealed that the GATase_6 and pribosyltran domains are conserved across fungi (Fig. 1A). Phylogenetic analysis of the Ade4 from different fungi revealed a close association of MoAde4 with the Ade4 protein of *Neurospora crassa* (Nc) and *Trichoderma reesei* (Tr) (Fig. 1B).

**Fig. 1.**
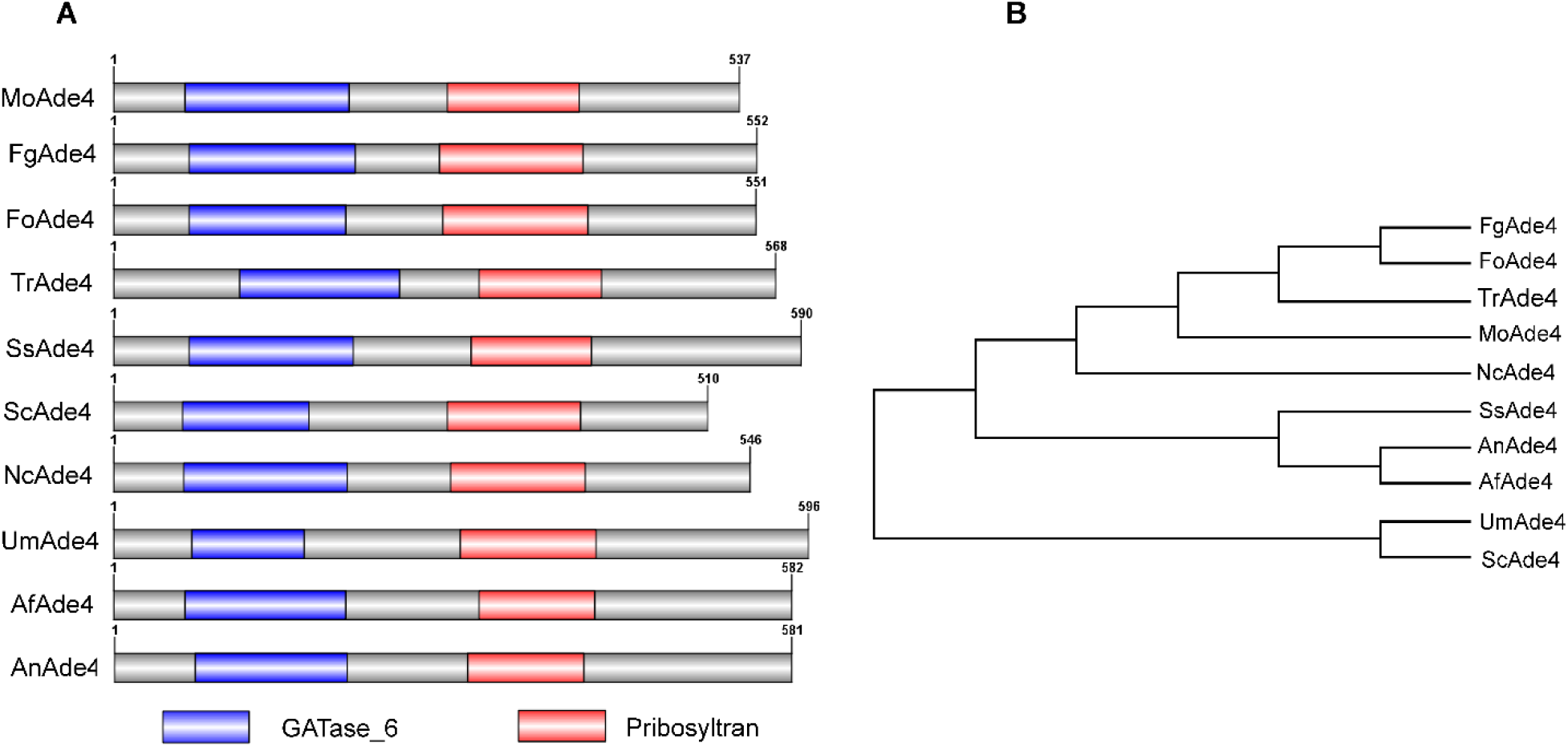
Domain architecture and phylogenetic analysis of MoAde4 protein from different fungal groups. (**A**) Domain architecture showing the conserved GATase and pribosyltran domains in different fungi. **(B**) Maximum Likelihood phylogenetic representation of MoAde4 protein in different fungi. The phylogenetic tree was generated using Mega software version 6 with a bootstrap method of 1000 replications.

### Expression of *MoADE4* gene at different developmental stages of *M. oryzae*

To gain insight into the likely roles of MoAde4 in *M. oryzae*, we firstly checked its expression at different development stages of the fungus. Using the mycelial stage for comparison, we found elevated expression of the *MoADE4* transcripts at the asexual, early-infection and late-infection stages of fungal development, with a 3.3-fold increase at the conidial stage and 2.3-fold, 1.8-fold 3.1-fold and 3.2-fold increases at 8, 24, 48, and 72 hours after inoculation respectively (Fig. 2). These results inferred that MoAde4 could be involved in both asexual development and infection in *M. oryzae*. To determine the role of MoAde4, we generated Δ*Moade4* deletion mutant by replacing the entire *MoADE4* gene from Guy11 with a hygromycin resistance gene using the split-marker homologous recombination method (Goswami, 2012). The deletion mutants were then confirmed by southern blot analysis (Fig S2).

**Fig. 2.**
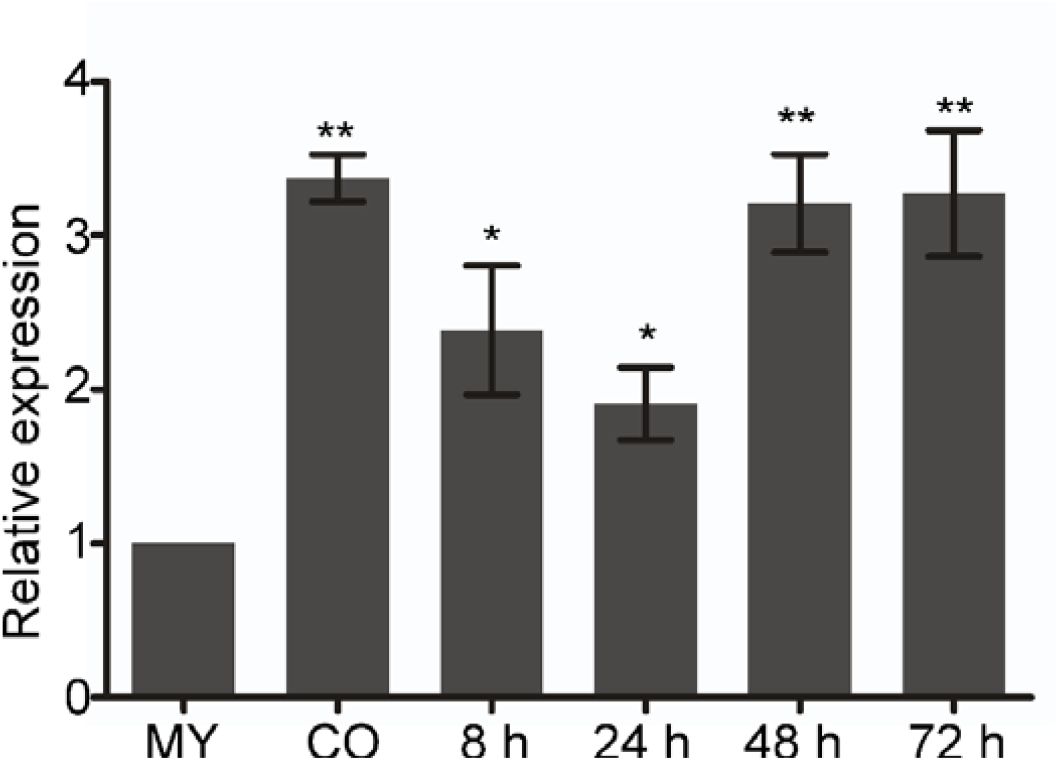
Expression patterns of *MoADE4* at different developmental stages. Expression was monitored at mycelia (MY), conidia (CO), and rice leaves inoculated with Guy11 conidia (5×10^4^ spores/mL) for 8, 24, 48, and 72 h. The *ACTIN* gene (MGG_03982) was used for normalization, the expression level of *MoADE4* at the mycelial stage was considered 1 for further comparisons. The qPCR results were obtained from three independent biological replications with three technical replicates. Error bars represent standard errors (*, *P* < 0.05; **, *P* < 0.01 by t-test).

### Loss of *MoADE4* resulted in reduced expression of other genes in *de novo* purine nucleotide biosynthetic pathway

Using *Saccharomyces cerevisae* genes that are known to catalyzed a series of the first ten reaction steps in the *de novo* purine biosynthesis pathway, we searched their remaining nine corresponding orthologs in *M. oryzae* genome and established, being encoded as follows, phosphoribosylaminoimidazole-succinocarboxamide synthase (MGG_12537) ADE1, phosphoribosylaminoimidazole carboxylase (MGG_01256) ADE2, C-1-tetrahydrofolate synthase (MGG_01014) ADE3, phosphoribosylformylglycinamidine cyclo-ligase (MGG_11343) ADE5, phosphoribosylformylglycinamidine synthase (MGG_11541) ADE7, phosphoribosylglycinamide formyltransferase (MGG_13813) ADE8, adenylosuccinate synthetase MGG_17000 (ADE12), adenylosuccinate lyase (MGG_03645) ADE13, and bifunctional purine biosynthesis protein (MGG_04435) ADE17. Next, we performed quantitative real-time PCR (qRT-PCR) analysis to establish the expression patterns of these nine genes upon deletion of *MoADE4*. Our results showed a down-regulation of all these genes in Δ*Moade4* (Fig. 3). These results therefore indicates, *MoADE4* that catalyzes the first step in purine biosynthetic pathway could be involved in positive regulation of the expression of the other genes in *de novo* purine nucleotide biosynthetic pathway

**Fig. 3.**
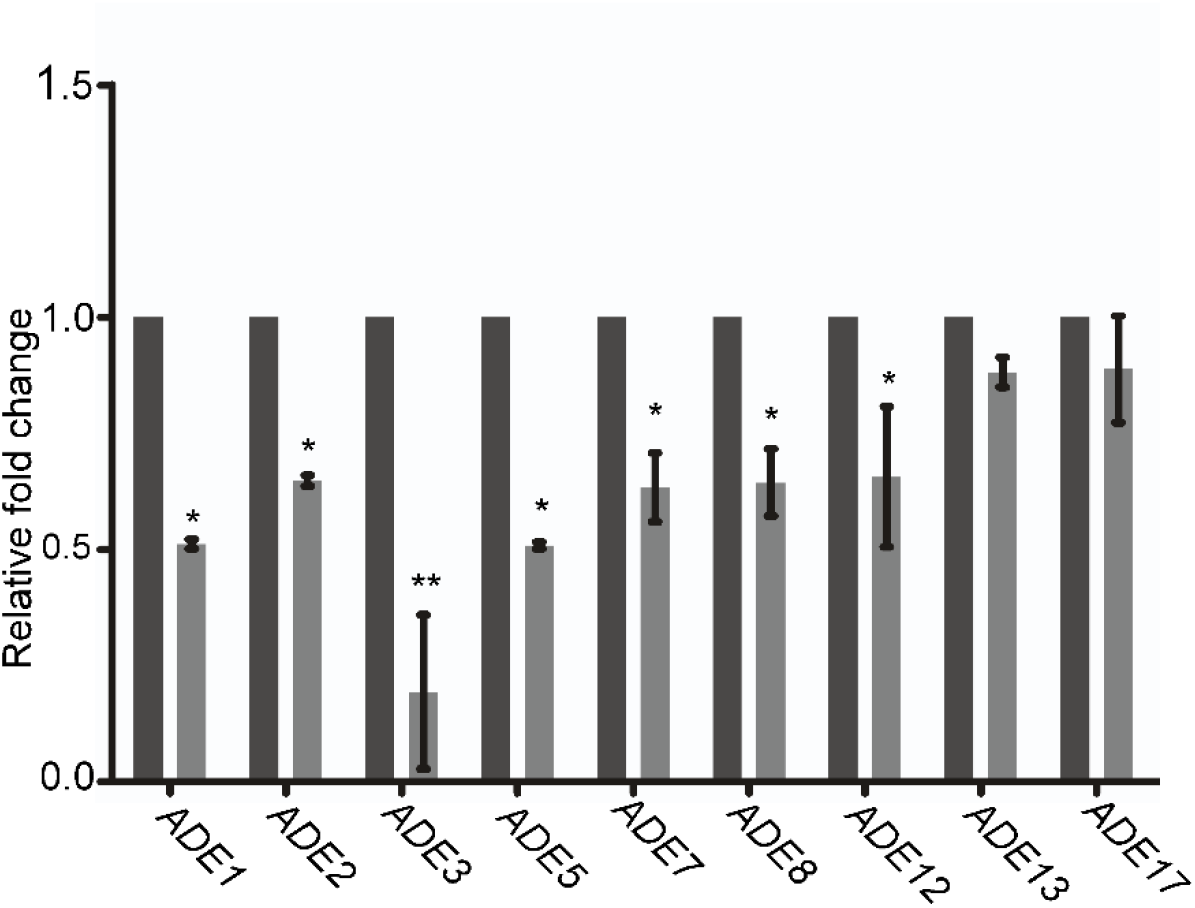
The expression levels of genes involved in *de novo* purine nucleotide biosynthesis pathway in the Δ*Moade4* mutant compared to the wild-type strain. The qPCR results were generated from three independent biological replications with three technical replicates. Error bars represent standard errors (*, *P* < 0.05; **, *P* < 0.01 by t-test).

### *MoADE4* is important for purine nucleotide biosynthesis

In *S. cerevisiae*, amidophosphoribosyltransferase (*ADE4*) acts as the first enzyme in the *de novo* purine biosynthesis pathway (Rolfes, 2006) To establish whether *MoADE4* gene is involved in purine nucleotide biosynthesis in *M. oryzae*, we initially cultured Guy11, Δ*Moade4* mutant, and complemented strains on complete medium (CM), straw decoction and corn medium (SDC), otmeal medium (OTM) and Minimum media (MM) and observed their growth. No significant difference was observed upon measuring the colony diameter of the Δ*Moade4*, WT, and the complemented strains after 10 days of incubation on CM and OTM (Fig. 4A). However, we noted a significant reduction in the development of aerial hyphae for Δ*Moade4* on OTM with hyphae becoming dark (Fig. 4A). The Δ*Moade4* deletion mutant could not sustain hyphal growth on MM (Fig. 4A). Previous study showed attenuated growth of Δ*Moade1* deletion mutant on MM with the recovery of growth occurring upon supplementation of MM medium with purine nucleotides adenine, adenosine and hypoxanthine (Fernandez et al., 2013). We therefore speculated that failure for Δ*Moade4* to grow on MM medium was due to deficiency in purine nucleotide. To address this, we cultured Δ*Moade4* in MM medium supplemented with purine nucleotides; adenine both (0.005 mM and 0. 01 Mm) and adenosine (0.01 mM, 0.1 mM, 1 mM). Our results showed that Δ*Moade4* strain was able to grow on MM media supplemented with adenine or adenosine (Fig. 4B and 4E), there was no difference in growth in terms of colony diameter and aerial hyphae development of Δ*Moade4* strain in both 0. 005 mM and 0. 01 mM adenine supplemented cultures. Development of Δ*Moade4* aerial hyphal on MM medium containing adenosine was dose dependent, with a high concentration (1 mM adenosine) showing well-developed developed aerial hyphae (Fig 4E). Purine nucleotide can be salvaged from hypoxanthine (Kappock et al., 2000), and previous study showed an operational salvage purine nucleotide pathway exists in rice blast fungus (Fernandez et al., 2013). To confirm whether deletion of *MoADE4* gene affected the normal functioning of the salvage pathway, we cultured Δ*Moade4* mutant on MM plates containing 0.01 mM hypoxanthine. Results showed rescued growth of Δ*Moade4* mutant on MM supplemented with hypoxanthine (Fig. 4C), thus indicating that loss of *MoADE4* impaired with purine nucleotide salvage pathway. Adenosine and other purine nucleotides were previously reported to be present in rice leaves (Sato et al., 2004), coupled with the fact growth of Δ*Moade1* occurred on MM medium prepared with rice leaves extract (Fernandez et al., 2013). We blended 100g of rice leaves, filtered the extract, and used it to prepare MM medium then cultured Δ*Moade4* on it. Results showed rescued growth of Δ*Moade4* strain on GMM medium prepared from rice extract (Fig. 4D). Taken together, these results demonstrates that *MoADE4* is important for both *de novo* and salvage purine nucleotide biosynthesis in rice blast fungus.

**Fig. 4.**
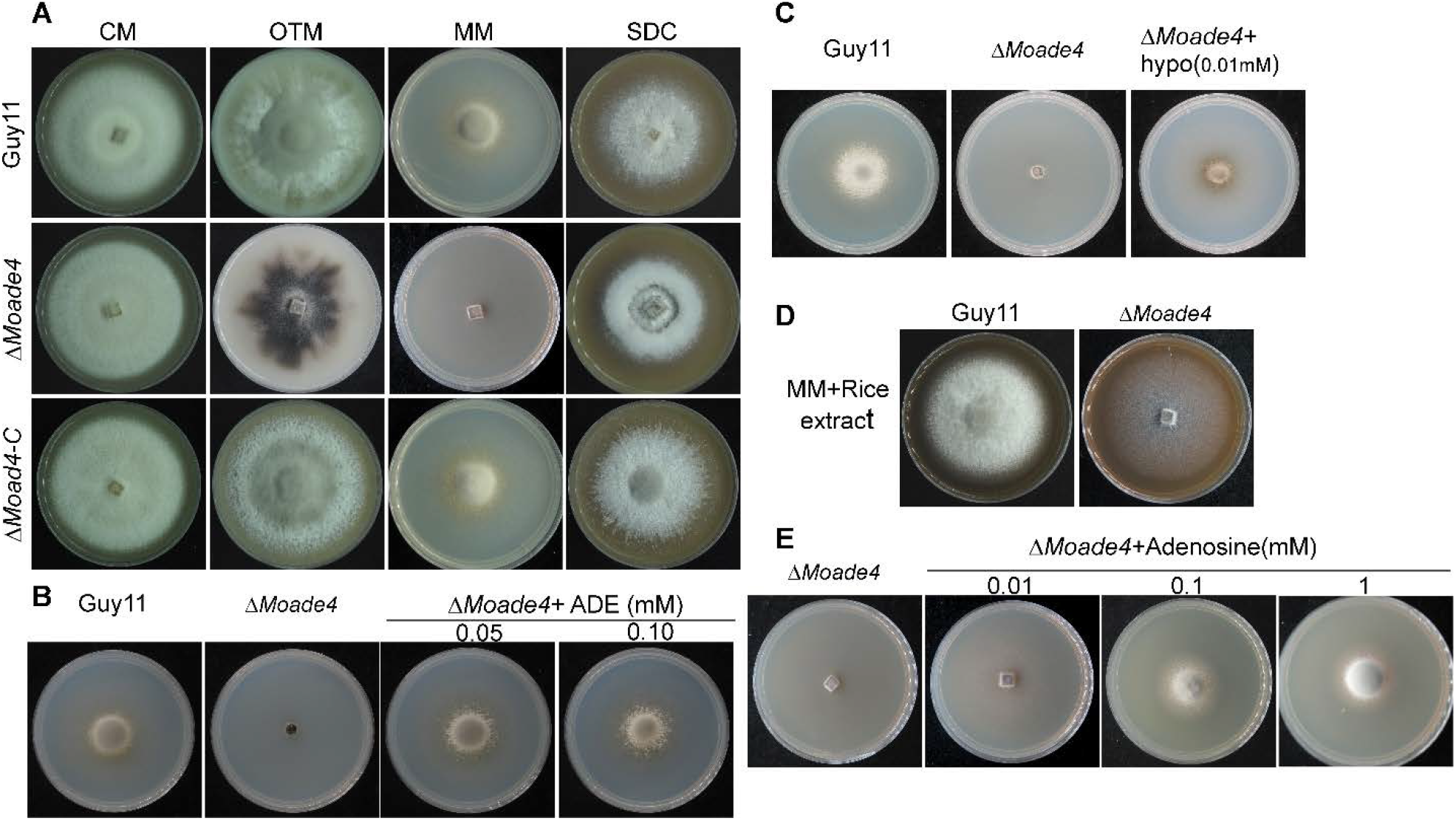
Effects of MoAde4 on hyphal growth. (**A**) Colony morphology and diameter of the Guy11, Δ*Moade4*, and complemented strain inoculated on complete medium (CM), otmeal medium (OTM), minimal media (MM) and straw decoction and corn medium (SDC). Plate images were photographed after 10 days. (**B**) Colony diameter of the Δ*Moade4* cultured on MM supplemented with 0.05mM and 0. 1mM adenine, photographs were taken after 8 days. (**C**) Plates showing colony diameter of Δ*Moade4*, growth being rescued on MM containing 0.01mM hypoxanthine, images were photographed after 8 days. (**D**) Photographs representation of Δ*Moade4* mutant showing recovery of growth on MM prepared from rice leaves extract. Images were taken after 8 days of inoculation. (**E**) Colony diameter of the Δ*Moade4* mutant cultured on MM supplemented with various concentrations (0.01 mM, 0.1 mM, 1. mM) adenosine. Photographs were taken 8 dpi

### MoAde4 contributes to asexual but not sexual reproduction in *M. oryzae*

Conidiogenesis is an important stage for the disease cycle of rice blast (Lee et al., 2006; Wilson & Talbot, 2009) To establish whether MoAde4 is involved in asexual reproduction in rice blast fungus. We incubated Guy11, Δ*Moade4*, and complemented strain in rice bran media for 10 days, then harvested and counted conidia for the three strains. We observed a significant reduction in aerial conidiophore development for the Δ*Moade4* mutant compared to WT Guy11 and the complemented strain (Fig. 5A). Concomitant with the reduced conidiophore development, we established a significant reduction in sporulation for the Δ*Moade4* deletion with 90% reduction in the number of conidia compared to the WT strain (Fig. 5B). Examination of the conidial structure showed no difference in conidia morphology between the Δ*Moade4* mutant and the reference strains (Fig. 5C). Next, we performed quantitative real-time PCR (qRT-PCR) analysis to examine the expression levels of genes important for conidiation and found reduced expression of *MoCOS1, MoCOM1, MoCON6, MoCON7*, and *MoHOX6* in Δ*Moade4* strain (Fig. 5D), thus indicating *MoADE4* positively regulates genes essential for conidiation. Furthermore, we performed mating experiment to establish the roles of *MoADE4* in sexual reproduction, the wild type Guy11 (MAT1-2), Δ*Moade4* and the complemented strain were crossed with a tester strain KA3 (MAT1-1) on oatmeal agar (OMA), thereafter perithecia production observed after three weeks. Perithecia formation occurred from the cross line of the tester strain KA3 with both Guy11, Δ*Moade4* and the complemented strain (Fig. S2), confirming that deletion of Moade4 did not affect ascospores production. In summary, these results indicates that *MoADE4* is required for asexual but not sexual reproduction in rice blast fungus.

**Fig. 5.**
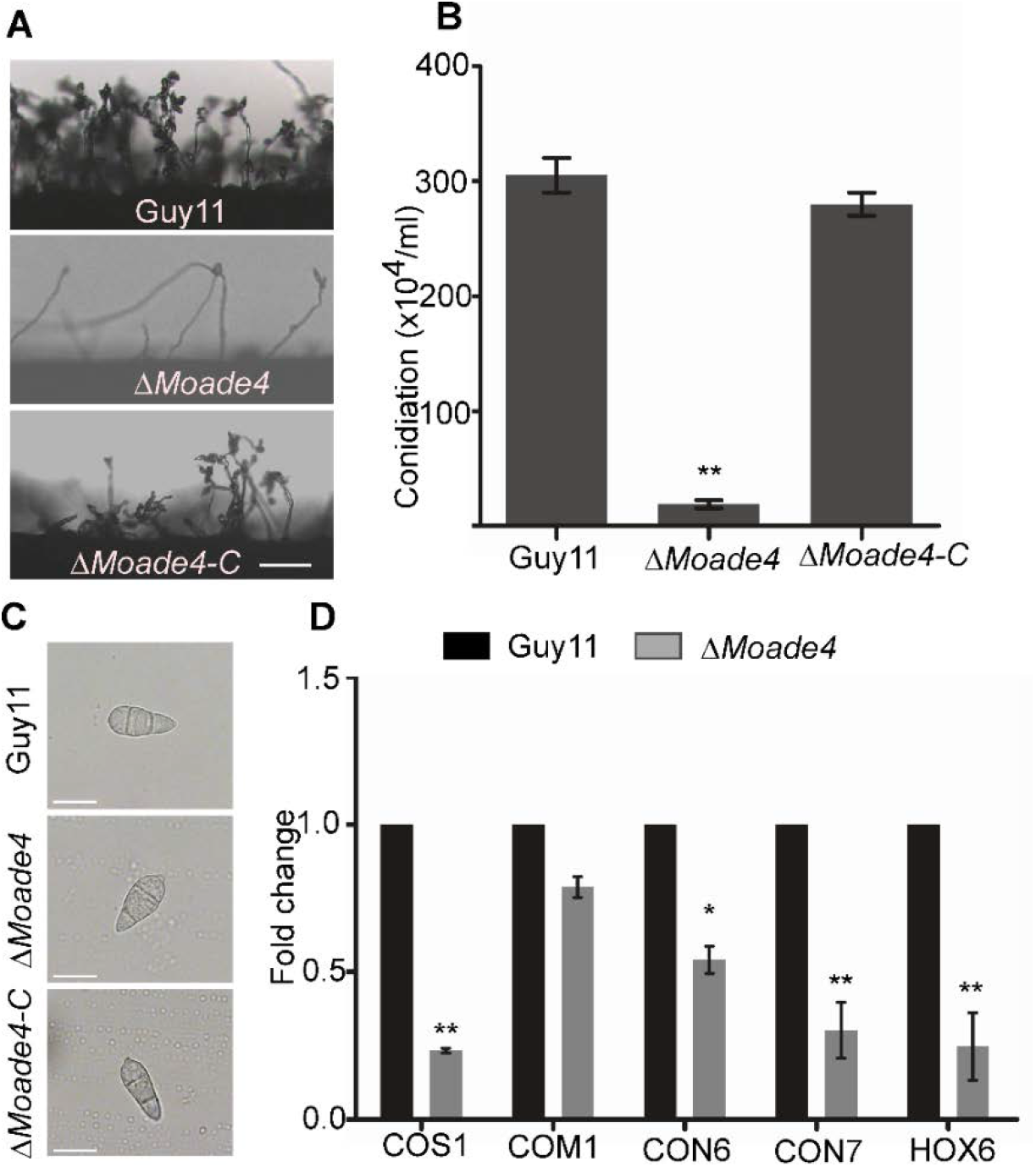
MoAde4 is required for conidiation of *M. oryzae*. (**A**) Development of conidia on conidiophores on rice bran observed under cover slips with a light microscope at 24 hpi after induction of conidiation. Bar= 10 μm (**B**). Conidiation ability of Guy11, Δ*Moade4* and the complemented strain on rice bran medium. The error bars represent standard errors from at least three independent replicates (**, *P* < 0.01 by t-test). (**C**) The expression levels of genes essential for conidiation in the Δ*Moade4* mutant compared to the Guy11 strain during vegetative growth. The qPCR results were obtained from three independent biological replications with three technical replicates. Error bars represent standard errors (*, *P* < 0.05; **, *P* < 0.01 by t-test)

### MoAde4 is essential for full virulence but not appressorium formation

To determine the effect of MoAde4 on the pathogenicity of rice blast fungus, conidia from Guy11, Δ*Moade4*, and the complemented strain were harvested for 10-day old rice bran medium, sprayed onto 3-week-old rice seedlings and disease progression monitored. No disease symptoms occurred on rice leaves infected with Δ*Moade4* conidia after 7days of inoculation, in contrast to leaves infected with Guy11 and complemented strains that displayed typical rice blast lesions (Fig. 6B). We also examined the infection profiles of the WT Guy11, the Δ*Moade4* mutant, and the complemented strain conidia on both intact and injured barley leaves, our results showed Δ*Moade4* was avirulent (Fig. 6C). Rice blast fungus asexual spores germinate into a specialized dome-shaped infectious structure called appressorium that facilitates the penetration of rice blast fungus (Saunders, Aves, & Talbot, 2010). Since the Δ*Moade4* mutant was nonpathogenic on rice and barley leaves, we speculated that the mutant might have defects in appressorial formation. To assess this possibility, we observed appressorial formation on inductive surfaces, our results showed that despite the Δ*Moade4* mutant being unable to cause disease both on rice and barley leaves, it was still able to produce appressoria on hydrophobic cover slips that were indistinguishable from the WT Guy11 strain and the complemented strain (Fig. 6A). These observations therefore, indicated that MoAde4 is essential for the fungal pathogenicity but not appressorium formation.

**Fig. 6.**
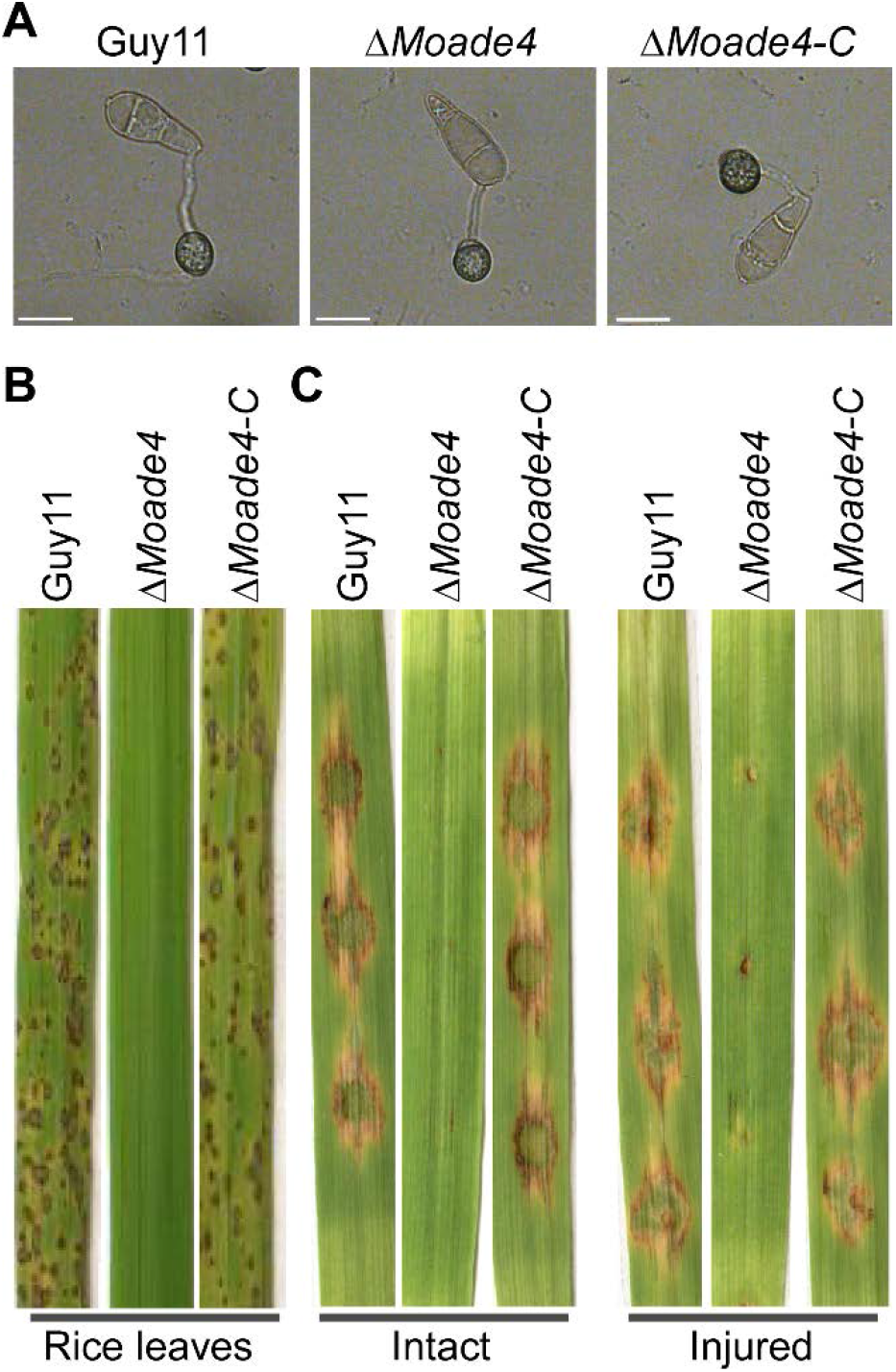
Appressorium formation and pathogenicity assay on rice and barley leaves (**A**) Appressoria of Guy11, the Δ*Moade4*, and the complemented strain were examined on hydrophobic cover slips using differential interference contrast (DIC) microscopy. Bar=25 μm. (**B**). Pathogenicity assays on rice (Oryza sativa cv. CO39). Rice leaves were sprayed with spores suspensions (5×10^4^ spores/mL) of Guy11, Δ*Moade4*, and the complemented strains and the leaves photographed 7 days post-inoculation. (**C**) Pathogenicity assays on barley. Conidial suspensions (10 µL, 5×10^4^ spores/mL) were inoculated onto barley leaves (intact or injured) for 7 days and then photographed.

### MoAde4 is required for appressorium penetration and successful colonization of host tissue

To account for the avirulent phenotype resulting from deletion of *MoADE4*, we inoculated rice sheaths with conidia from Guy11, Δ*Moade4* mutant, and the complemented strain and observed the appressorium formation, appressorium penetration, and the infectious hyphal growth. In contrast with Guy11 and the complementation strains whose appressorium had penetrated at 30 hours post-inoculation (hpi), the Δ*Moade4* appressoria were completely unable to penetrate the rice sheath while only approximately 50% penetrated at 48 hpi and 72 hpi. (Fig. 7A, and 7B). Examination of the invasive hyphae growth showed that, unlike Guy11 and the complemented strain, whose invasive hyphae were evident 30 hpi and had spread to the secondary cells at 48hpi. The Δ*Moade4* had not formed invasive hyphae at 30 hpi only forming at 48hpi (Fig. 7A), and they were restricted to the primary cells and could not spread to the adjacent neighboring cells (Fig. 7C). These findings suggest MoAde4 plays an important role in appressorium penetration, host colonization and thus causing the pathogenicity defect of *M. oryzae*.

**Fig. 7.**
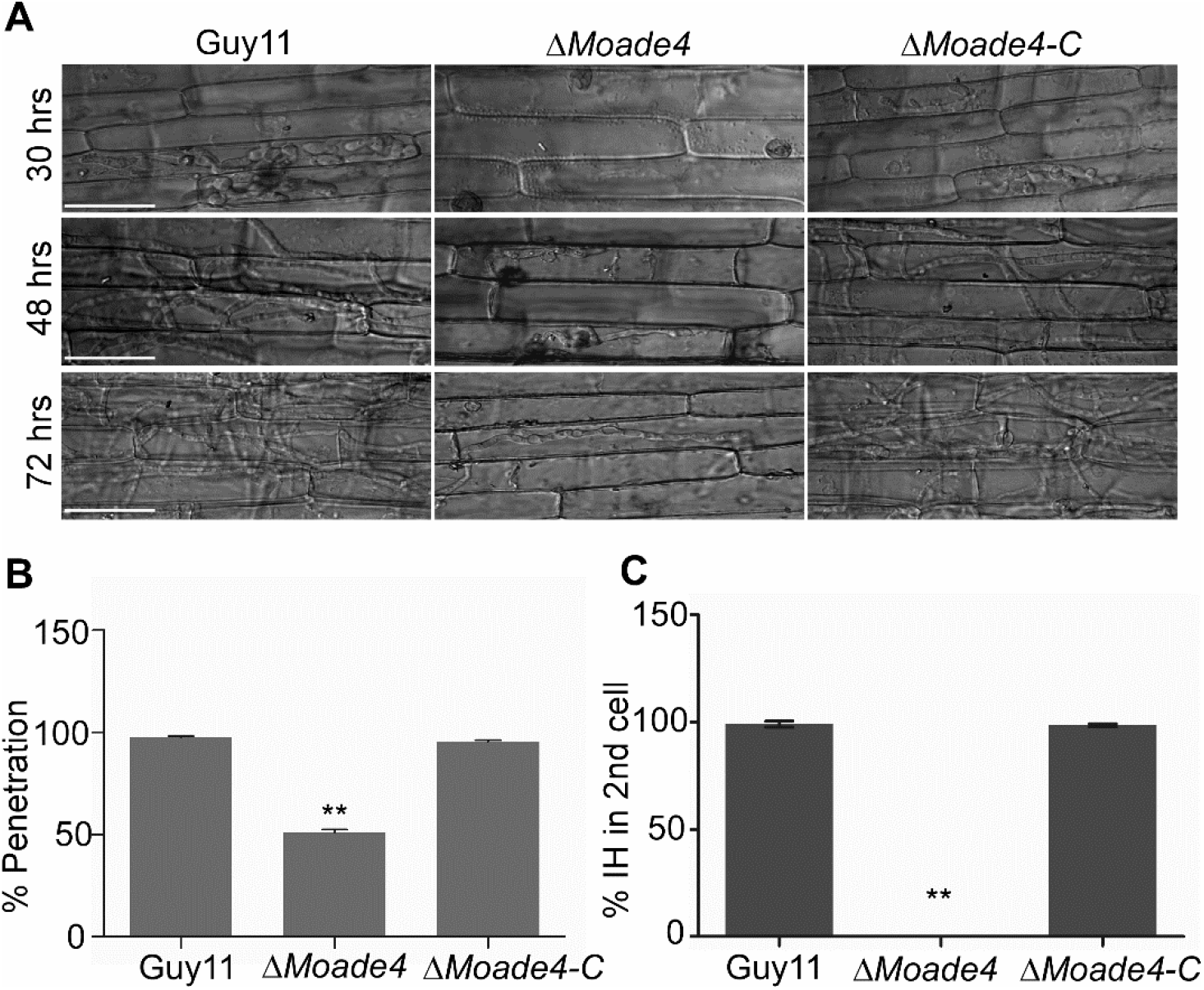
MoAde4 is important for appressorium penetration and IH colonization. (**A**) The infection structures formed on rice epidermal cells in response to inoculation with conidia of the Guy11, Δ*Moade4*, and complemented strain in rice sheath at 30, 48, and 72 h. Bar= 20 µm. **(B**) Statistical representation of the percentage of appressorium penetration into rice cells. The error bars represent standard errors from at least three independent replicates (**, *P* < 0.01 by t-test). (**C**) Statistics showing the percentage of invasive hyphae that colonize the rice sheath secondary cells, the Δ*Moade4* mutant invasive hyphae failed to extend the secondary cells. The error bars represent standard errors from three independent replicates (**, *P* < 0.01 by t-test).

### MoAde4 is essential for appressorium glycogen mobilization and turgor pressure build up

In rice blast fungus, glycogen reserves has been reported to be abundant in conidia and is consumed rapidly during conidial germination (Bourett & Howard, 1990; Thines, Weber, & Talbot, 2000). We reasoned out that the penetration defect exhibited by Δ*Moade4* might be caused impaired mobilization of glycogen during conidial germination. To test this phenomenon, we used KI/I2 to stain conidia and appressoria of the wild type Guy11, Δ*Moade4* mutant and the complement strain. Results showed abundant distribution of KI/I2-stained glycogens of the Guy11, Δ*Moade4* mutant and the complement strain during germination of conidia on hydrophobic surfaces from 0 to 4 hpi (Fig. 8A and 8B). However, glycogen mobilization from the Δ*Moade4* conidia to appressoria was remarkably retarded from 8 to 24 hpi compared to the wild type and the complemented strain (Fig. 8A and 8C). In addition to glycogen, host tissue penetration and the development of invasive hyphae require sufficient build up hydrostatic pressure within the appressorium. Glycerol assays have been used to assess appressorial turgor (Howard, Ferrari, Roach, & Money, 1991; Z.-Y. Wang et al., 2005). Our cytorrhysis (appressoria collapse) assay indicated that the turgor pressure of the Δ*Moade4* mutant was much lower than that of the wild-type and the complemented strain as evident by increased appressoria collapse upon treating with 1M, 2M, and 3M glycerol concentrations (Fig. 8D and 8E). Taken together, these results indicates *MoADE4* plays important roles in conidium to appressorium glycogen mobilization and turgor pressure build-up thus, contributing immensely in appressorium penetration during in planta infection in rice blast fungus.

**Fig. 8.**
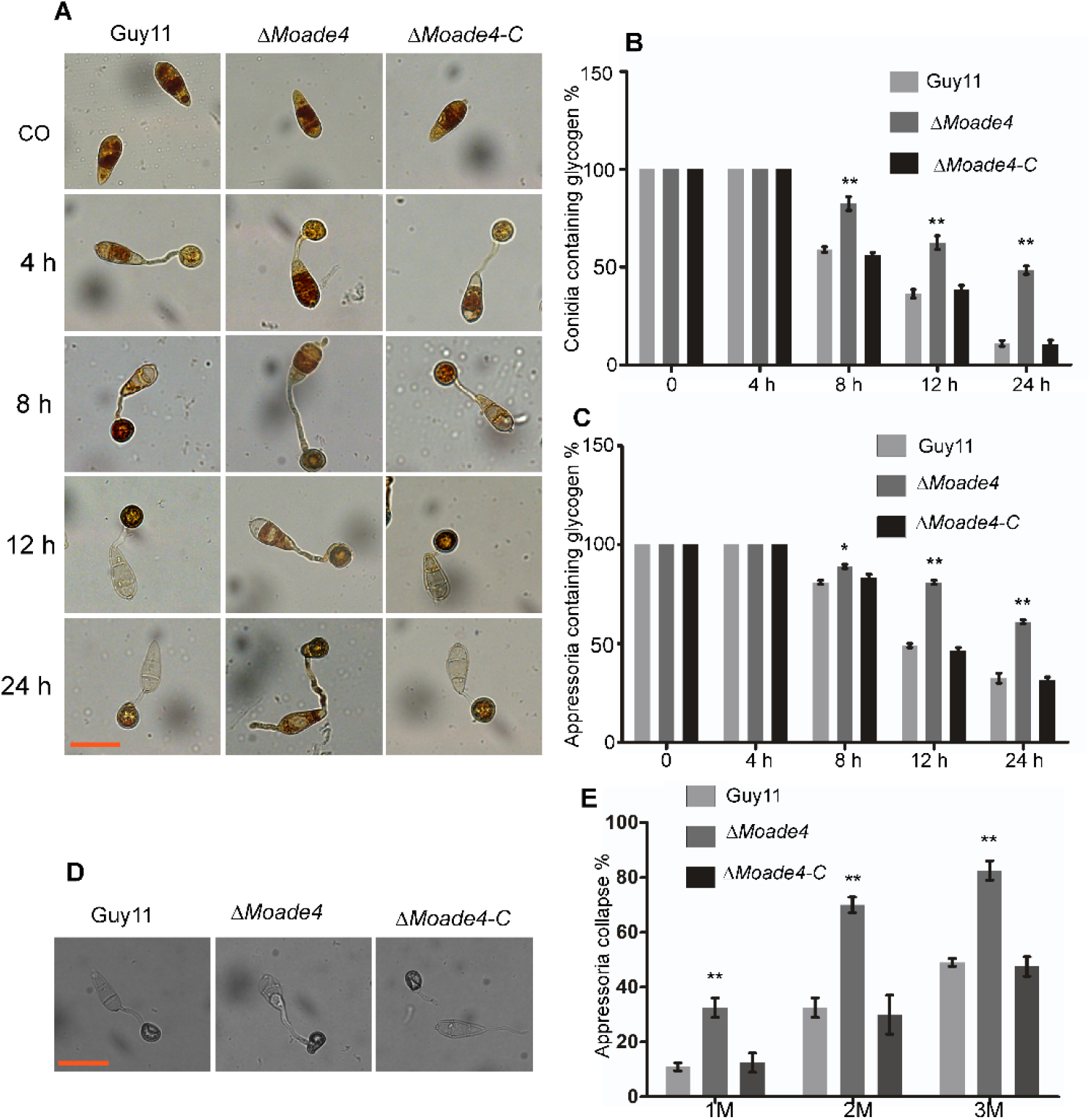
Deletion of *MoADE4* leads to delayed mobilization of glycogens from conidia to appressoria and lowered appressorium turgor. (**A**) Conidia and appressoria of the wild-type Guy11, Δ*Moade4* and the complemented strain that germinated on hydrophobic coverslip. Samples were stained with KI/I2 solution at 0, 4, 8, 12, and 24 hpi. Bar = 10 µm. (**B**) Bar graph showing the percentage of the Guy11, Δ*Moade4* and the complemented strain conidia containing glycogens, 100 conidia were counted each time. Error bars represent standard errors with three independent replicates (**, *P* < 0.01 by t-test). (**C**) Graphical representation of the percentage appressoria of the Guy11, Δ*Moade4* and the complemented strain having glycogens, 100 appressoria were measured. The error bars represent standard errors with three independent replicates (*, *P* < 0.05; **, *P* < 0.01 by t-test). (**D**) Appressorium of the Guy11, Δ*Moade4* and the complemented strain that collapsed after being incubated with glycerol Bar = 10 µm. (**E**) Bar graph showing Statistical analysis of collapsed appressoria (24 h) of the Guy11, Δ*Moade4* and the complemented strain after incubation for 15 min in 1M, 2M and 3M glycerol solutions. 100 appressoria were counted per experiment. The error bars represent standard errors of appressoria that had collapsed with three independent replications (**, *P* < 0.01 by t-test).

### Purine transporters genes in Δ*Moade4* and rice pathogenesis genes were down-regulated at 48h during in *planta* infection

Plants respond to invading pathogens by induction of defense programs required to neutralize the pathogens, this has to be avoided by pathogens for a compatible interaction to occur (Chi, Park, Kim, & Lee, 2009; Mentlak et al., 2012). We therefore speculated that the restricted invasive hyphal growth by Δ*Moade4* mutant in rice cells (Fig. 7A and 7C), could be caused by increased predisposition to host defenses and nutrient deficiency. To test this, we firstly analyzed the expression of rice pathogenesis-related (PR) genes by qRT-PCR to determine if Δ*Moade4* strain elicited stronger rice defense responses relative to Guy11 during infection. Our results showed reduced expression of rice -PR genes *PR1a, PR1b*, and *PBZ1* in rice challenged with Δ*Moade4* strain compared to the WT Guy11 (Fig. 9A) and contradicts with the previous study where the expression of PR genes, PBZ1 and PR1, were shown to be more highly induced by Δ*des1* infection than by WT (Chi et al., 2009).

**Fig. 9.**
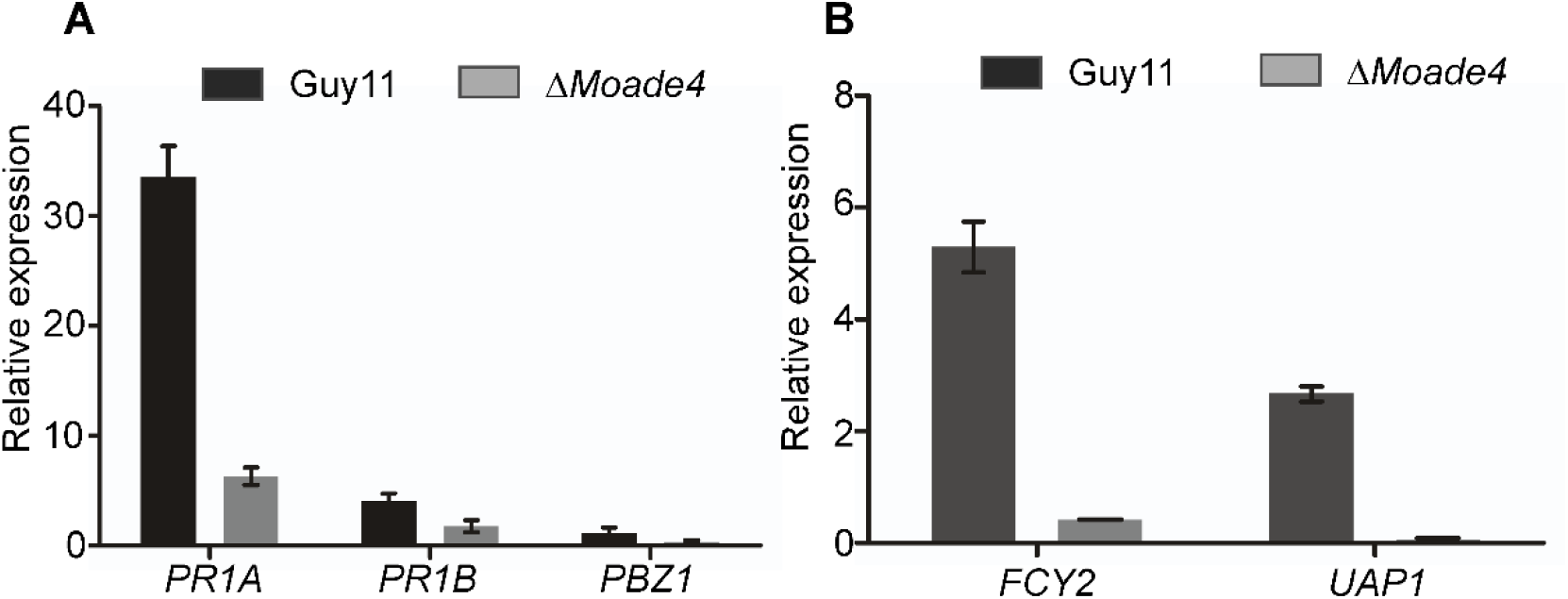
Δ*Moade4* mutant failed to elicit strong plant defense responses compared to Guy11 and purine transporter genes are repressed in rice infected with Δ*Moade4*. (**A**) The expression of the rice pathogenesis-related genes *PR1A, PR1B*, and *PBZ1* in rice leaves challenged with Guy11 and Δ*Moade4*. The qPCR results were obtained from three independent biological replications with three technical replicates, error bars represent standard deviation. The expression of rice OsACT2 was used as a control. (**B**) Expression of purine transporter genes *FCY2* and *UAP1* during infection and normalized against the expression of MoAct. The data were generated from three independent biological replicates with three technical replicates, error bars represent standard deviation

Plant fungi acquire nutrients from their host that facilitate their effective interactions, but our findings show that the poor development of Δ*Moade4* invasive hyphae during infections could be a result of insufficient purine amounts. A previous study showed the availability of purine adenosine in rice leaves (Sato, Soga, Nishioka, & Tomita, 2004). Coupled with the fact Δ*Moade4* was able to grow on both adenosine and MM plates made of rice extract (Fig. 4D and 4E), we expected Δ*Moade4* to acquire purine from rice leaves required to facilitate its invasive hyphae colonization in rice cells. However, the growth of the invasive hyphae was curtailed, which prompted us to speculate a possible defect in purine uptake from rice leaves. To address this, we analyzed the expression of two genes that encode purine transporter genes, purine-cytosine permease *(FCY2)* (Gournas, Oestreicher, Amillis, Diallinas, & Scazzocchio, 2011), and uric acid-xanthine permease *(UAP1)* (Ferreira, Chevallier, Paumard, Napias, & Brèthes, 1999). Results showed that these genes were repressed during infection in rice leaves challenged with Δ*Moade4*, and were up-regulated in rice infected with Guy11 (Fig. 9B). The repression of purine transporters genes in Δ*Moade4* during infections possibly explains why Δ*Moade4* could not get purines from rice leaves. Based on these results, we concluded that the retarded invasive hyphae growth for Δ*Moade4* in rice cells was not a result of host immunity but due to purine deficiency.

### Exogenous adenine partially restored defects in conidiation, invasive hyphal colonization and pathogenicity of Δ*Moade4*

Since Δ*Moade4* mutant exhibited adenine auxotroph on MM (Fig. 4C), we postulated that the defects in sporulation, invasive hyphal growth and colonization and pathogenicity were as a result of a dysfunctional *de novo* purine biosynthetic pathway. To test this idea, we inoculated the Δ*Moade4* mutant in rice bran media augmented with 0.01 mM and 0.005 mM adenine. The spores derived from 0.01 mM adenine-supplemented cultures were inoculated in rice sheath for observation of invasive hyphal development and sprayed on rice leaves then applied on barley leaves for pathogenicity assays. Our results showed a partial restoration in conidiation as the number of spores doubled when compared to those produced on rice bran culture without adenine (Fig. 10A). In addition, pathogenicity assays showed an improved virulence of Δ*Moade4* spores derived from 0.01 adenine cultures both on barley and rice leaves (Fig. 10B and 10C). Lastly, examination of the invasive hyphal development revealed exogenous adenine was able to rescue the defects in invasive hyphae proliferation from the primary cells to the adjacent rice cells (Fig. 10D). These results therefore, confirms that *de novo* purine nucleotide biosynthesis is important for conidiation, invasive hyphal colonization and pathogenesis of *M. oryzae*.

**Fig. 10.**
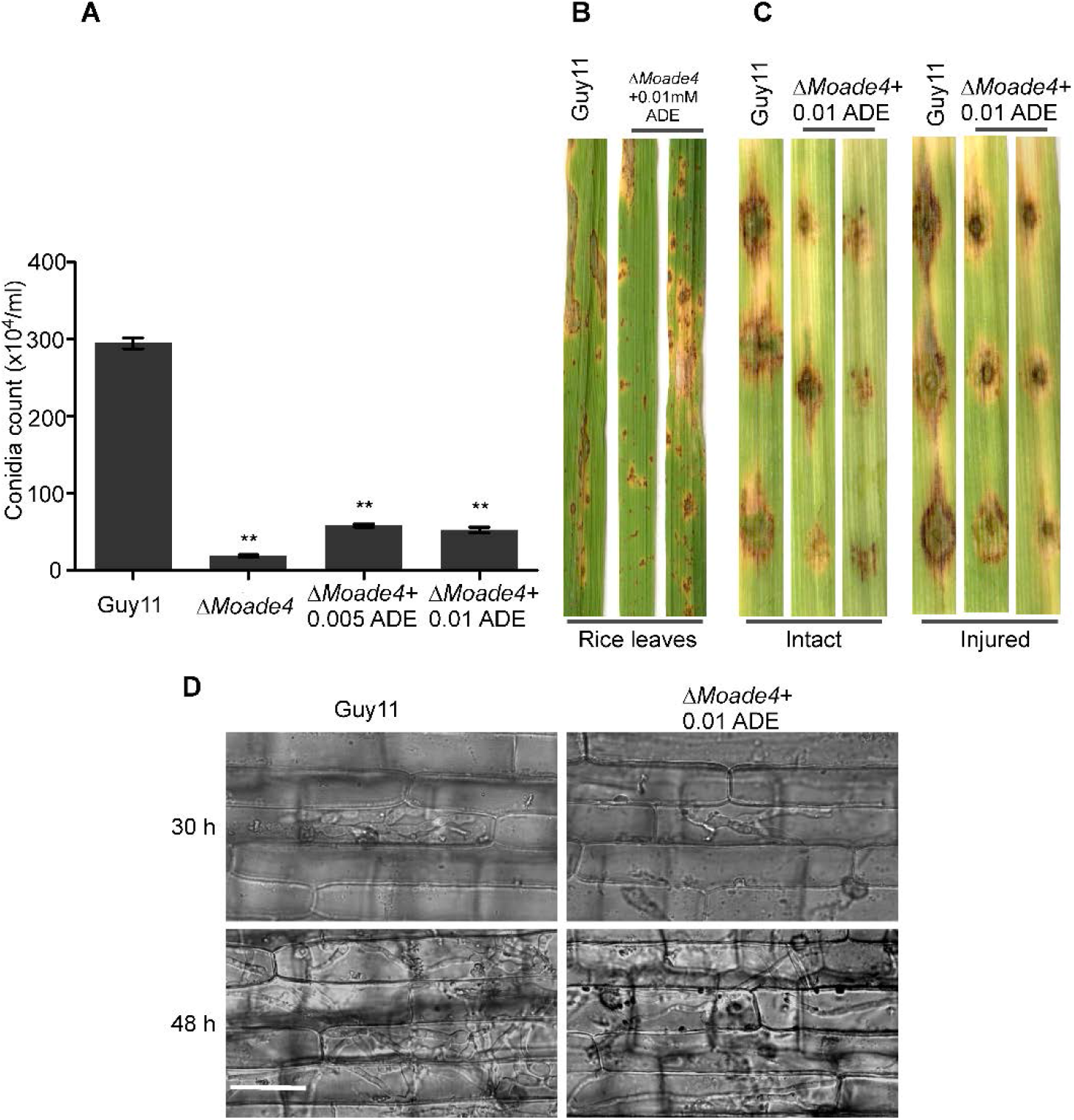
Exogenous adenine partially restores conidiation, pathogenicity and invasive hyphal colonization defects exhibited by Δ*Moade4* strain. (**A**) Statistics showing the conidiation characteristics of Δ*Moade4* compared with Guy11 strain, Δ*Moade4* was cultured in rice bran media supplemented with 0.005 mM and 0.01 mM adenine. The error bars represent standard errors from at least three independent replicates (**, *P* < 0.01 by t-test). (**B** and **C**) Pathogenicity of the strains on rice seedlings and barley leaves respectively. Conidia (1×10^5^ /mL) were harvested from 0.01 mM adenine-supplemented cultures, 0.01 mM adenine was added to the conidial suspensions, and then sprayed onto 3-week-old rice seedlings, some dropped onto 10-day-old barley leaves, and then photographed at 7 dpi. **D** Infection structures formed on rice cells in response to inoculation with conidia of the WT, and Δ*Moade4* spores derived from 0.01 adenine cultures at 36, and 48 hpi.

### Subcellular localization of the MoAde4 protein

To establish the cellular component to which the MoAde4 protein is located, the MoAde4-GFP fusion construct and a plasmid containing a neomycin resistance gene were transformed into Δ*Moade4* mutant protoplasts. MoAde4-GFP was targeted to the cytoplasm in the growing mycelia, conidia, appressorium and invasive hyphae during infection (Fig. 11), thus showing that MoAde4 is localized in the cytoplasm.

**Fig. 11.**
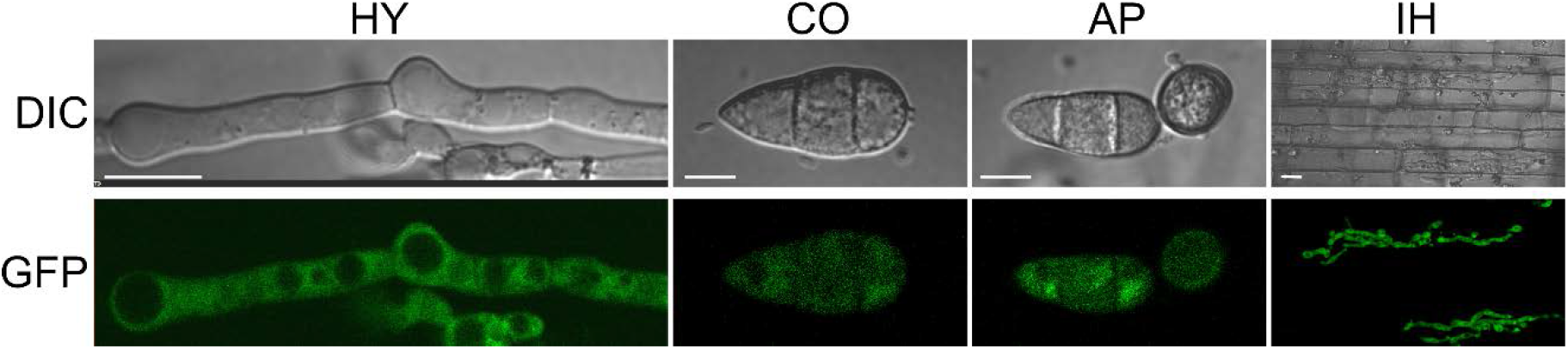
Sub-cellular localization of MoAde4 in *M. oryzae*. MoAde4-GFP is localized in the cytoplasm in growing hyphae (HY), conidia (CO), appressorium (AP), and invasive hyphae (IH) during infection. Scale bar =10 µm

## Discussion

Biosynthesis of purine nucleotide by *de novo* and salvage pathways has been shown to be important for growth, development, and pathogenicity in various microorganisms (Kirsch & Whitney, 1991; Morrow et al., 2012; Rodriguez-Suarez et al., 2007), thus enzyme of these pathway serves as an important antifungal target. In this study, we identified and characterized MoAde4 a homolog of *S. cerevisiae* Ade4. Phylogenetic analysis revealed that MoAde4 is 537-amino acid protein consisting of the GATase_6 and pribosyltran domains. Additional Ade4 domain prediction for the other fungi revealed that these two domains were conserved, suggesting a likelihood of similar functions. Deletion of *MoADE4* did not result in any significant difference in the growth of Δ*Moade4* mutant on nutrient-rich CM media and SDC while the mutant exhibited dark pigmentation on OTM. However, the hyphal growth of Δ*Moade4* mutant was attenuated on MM and was restored on MM supplemented with adenine, adenosine, and hypoxanthine. These results were similar to those for the study in *C. neoformans*, where purine bases were required for the growth of mutants in the purine biosynthesis pathway (Morrow et al., 2012). Additional growth of Δ*Moade4* mutant was also evident on GMM plates prepared with rice leaves extract, an explanation for this is because rice leaves contain purine nucleotide(Sato et al., 2004).

Conidiogenesis is an important step in the disease cycle of rice blast (Lee et al., 2006). Similar to reduced conidiation observed on the deletion of phosphoribosylamidotransferase in sclerotial mycoparasite *Coniothyrium minitans* (Qin et al., 2011), conidia quantification assay revealed Δ*Moade4* deletion mutant was compromised in terms of asexual reproduction, with a 90% reduction in sporulation relative to that of the WT strain. Analysis of the expression of genes involved in conidiation (*COS1, COM1, CON6, CON7*, and *HOX6*) revealed a down-regulation of these genes in the Δ*Moade4* mutant, suggesting that *MoADE4* gene likely regulates the expression of these of these conidiation related genes. These results were similar to previous findings where a defect in asexual reproduction exhibited by Δ*Moppe1* mutant was linked to the down-regulation of genes important for conidiation (Qian et al., 2018). Exogenous adenine partially rescued spore production in the Δ*Moade4* deletion mutant, however, the conidia number was still lower than that of the WT strain. The reduced number of conidia produced by Δ*Moade4* mutant and partial restoration of conidiation when adenine was added in sporulation media demonstrates that *de novo* purine nucleotide biosynthesis mediated by *MoADE4* is indispensable for conidiogenesis in the rice blast fungus.

Appressoria are critical for *M. oryzae* penetration of host cells (Howard et al., 1991; Saunders et al., 2010). Deletion of *MoADE4* resulted in almost of 50% its appressoria being unable to penetrate the rice cells, analysis of conidia to appressorium glycogen mobilization and turgor pressure which are key requirements for appressorium function, showed delayed glycogen mobilization and reduced turgor pressure in the appressoria of the Δ*Moade4* deletion mutant. These findings echoed previous studies that demonstrated impaired appressoria penetration in rice cells were as a result of the defects in mobilization of glycogen from conidia to appressorium for Δ*Moglt1* (Zhou et al., 2018) and turgor pressure for the Δ*Molys2* (Y. Chen et al., 2014). Examination of invasive hyphae showed a delayed infectious hyphae growth which occurred at 48 hpi and they were restricted to the primary host cells.

The Δ*Moade4* mutant was avirulent on rice and both intact and injured barley leaves. This is similar to what was observed in the human fungal pathogen *Candida albicans*, where adenine deficiency abolishes virulence in a murine model (Jiang et al., 2010). The virulence defect displayed by Δ*Moade4* mutant was caused by restricted invasive hyphae growth within the primary cells. In rice blast fungus, pathogenicity defects of mutants resulting from deletions of genes important for amino acid biosynthesis such as *LYS2, CPA2, MET13*, and *ARG1/5/6/7* could be rescued upon addition exogenous amino acids (Y. Chen et al., 2014; Liu et al., 2016; Yan et al., 2013; Y. Zhang et al., 2015). Just like in these mutants, the virulence defects of Δ*Moade4* mutant was restored by exogenous adenine, thus, demonstrating that *de novo* purine nucleotide biosynthesis mediated by MoAde4 is crucial for fungal virulence in *M. oryzae*. The expression of PR genes upon fungal infection is one of the parameters for studying the host defense response. Studies on the host defense response to rice blast fungus through the expression of PR genes have been reported (Guo et al., 2010; Liu et al., 2016). In this study, we established that Δ*Moade4* mutant could not induce a strong host immune response as evident by lower expression of PR genes (*PR1A, PR1B, PPZ1*) upon rice infection relative to the WT strain, and contradicts the previous study where *Moades1* mutant was nonpathogenic due to elevated expression of PR genes in rice challenged with Δ*Moades1* mutant compared to the WT strain (Chi et al., 2009). The relative low induction of host immunity by Δ*Moade4* demonstrates that the host immunity was not the main reason for the Δ*Moade4* restricted invasive hyphal growth during infection but instead purine nucleotide deficiency. Although we have not shown that the addition of exogenous purine could restore the invasive hyphae colonization in Δ*Moade4* mutant during infection, the partial restoration of pathogenicity defect by Δ*Moade4* spores obtained from 0.005mM adenine supplemented culture on rice and barley leaves clearly shows an improved invasive hyphae colonization. Pathogens acquire nutrient during interactions (Glazebrook, 2005). Previous study on obligate biotrophic plant pathogen *Erysiphe graminis* showed it was able to acquire purine from its host during infections (Butters, Burrell, & Hollomon, 1985). We expected a similar scenario for Δ*Moade4* mutant to acquire purine nucleotide from rice leaves during infection that will help facilitate host invasive hyphae colonization, but this was not the case, a possible explanation for the failure to acquire purine from rice could be the repression of purine transporter genes *(FCY2)* and *(UAP1)* in Δ*Moade4* required for uptake purines as evident by qPCR analysis.

In summary, our characterization of the biological functions of MoAde4 in *M. oryzae* revealed that *de novo* purine nucleotide biosynthesis mediated MoAde4is essential for appressorium-mediated penetration and infectious hyphae colonization of plant tissues, thereby influencing pathogenicity of *M. oryzae*. These findings provide further evidence for the understanding of the pathogenic mechanisms of phytopathogens and serves are an important target for antifungal development.

## Material and methods

### Fungal strains and culture conditions

The wild type (WT) Guy11 and mutant strains of *M. oryzae* were cultured at 25°C using complete media (CM: 0.6% yeast extract, 0.6% casein hydrolysate, 1% sucrose, 1.5% agar) as described (J. Chen et al., 2008). Other media used in this study included minimal media (MM: 6 g of NaNO3, 0.52 g of KCl, 0.52 g of MgSO4, 1.52 g of KH2PO4, 10 g of glucose, and 15 g of agar in 1 L of double-distilled water), straw decoction and corn media (SDC: 100 g of rice straw, 40 g of corn flour, and 15 g of agar in 1 L of double-distilled water) and oatmeal agar media (OTM: 50 g of oatmeal and 15 g of agar in 1 L of double-distilled water).

Samples for genomic DNA extraction, total RNA and protoplast preparation were cultured in liquid CM in an orbital shaker at 110 rpm for 3 days.

To induce conidiation, strains were cultured on rice bran agar (2% rice bran, 1.5% agar; pH 6.5) for 10 days at 28°C in the dark followed by 3 days of continuous light illumination. Conidia were collected in 5 ML of distilled water, filtered using three-layer lens paper, and counted with a hemocytometer under a light microscope.

### Target gene deletion and complementation in *M. oryzae*

Split-marker approach (Goswami, 2012) was adopted in the generation of Δ*Moade4* deletion mutant in *M. oryzae* by replacing the entire *MoADE4* gene with the hygromycin gene (*HPH*). The upstream and downstream flanking sequences of the *MoADE4* were amplified from *M. oryzae* genomic DNA with primer pairs (ADE4F1/ ADE4F2 and ADE4F3/ ADE4F4) respectively. The resultant PCR products were used to generate split makers by ligating with hygromycin fragments amplified from pCX62 using primer pairs HYG/F(F)/HY/R(R) and YG/F(F)/HYG/R(R) (Supplementary Table S2).

Fungal protoplast preparation and polyethylene glycol (PEG)-based transformation of *M. oryzae* was performed as described previously(Wendland, 2003; C. Zhang et al., 2016) Putative hygromycin and neomycin-resistant candidates transformants were screened on TB3 media supplemented with 250 μg/mL hygromycin B (Roche Applied Science) and 200 μg/mL G418 (Invitrogen) and the mutants verified by Southern blotting analysis.

For complementation, the 4.2-kb PCR product containing about 2.4-kb native promoter and the full-length *MoADE4* gene coding region were first amplified using primers ADE4C1(F)/ADE4C2(R) and cloned into pKNTG. After verification of the sequence, the construct was transformed into the protoplast of the Δ*Moade4* mutant. The complemented strains were screened by neomycin resistance and observations of GFP fluorescence.

### Appressorium formation, penetration and infection assays

Conidia collected from 10 days old rice-bran culture were adjusted to (5×10^4^ spores/mL) using sterilized double-distilled water with 0.02% (v/v) Tween-20 solution. For appressorium formation assays, 20 µL of conidial suspension was added to an artificial hydrophobic coverslip and incubated in darkness at 28°C. Appressorium formation was then examined at 4 h, and 8 h time intervals.

Conidia germination and appressorium formation on inductive surfaces were measured as described before (Qi et al., 2012)

Rice infection was performed by spraying 3-week-old rice (*Oryzae sativa* cv. CO39) seedlings with a conidial suspension (5×10^4^ spores/mL), the infected plants were incubated in a humid chamber at 28°C for 24 h in darkness and later transferred to a12-h photoperiod chamber. Leaves were then imaged 7 days after infection. For the barley infection assay, 10 µL of conidial suspension (5×10^4^ spores/mL) were dropped on barley leaves and incubated at 28°C for 24 h in darkness. Later, they were transferred to light conditions and imaged after 7 days.

For the rice sheath penetration and invasive hyphae growth assay, 100 µL of conidial suspension (5×10^4^ spores/mL) was inoculated into the inner rice sheath cuticle cells. The plants were then incubated for 30 h 48 h and 72 h at 28°C under humid conditions, and the leaf sheaths were examined using a microscope as described before (Zhang et al., 2011).

### Nucleic acid manipulation, southern blotting analysis, and qRT-PCR

The cetyltrimethylammonium bromide (CTAB) method was used for the extraction of genomic DNA from the WT strain Guy11 and Δ*Moade4* deletion mutants (Y. Wang et al., 2017).

For southern blot assays, probes for the *MoADE4* gene and the hygromycin (HPH) gene were amplified with primer pairs ADE4 ORF1/ADE4 ORF2 and Hph F/ HphR respectively (Table S2). Probe labeling, hybridization, and detection were performed with a DIG High Prime DNA Labeling and Detection Starter Kit (Roche Applied Science, Penzberg, Germany).

For measuring the relative abundance of MoAde4 transcripts during various stages of fungal development. RNA samples were obtained from Guy11 mycelia cultured in liquid CM in an orbital shaker at 110 rpm for 3 days, conidia, and from plants infected with Guy11 conidia (1 × 10^8^ spores/mL) for 8, 24, 48, and 72 h.

To evaluate the transcript level of PR genes, RNA samples were from the rice leaves inoculated with Guy11, and the Δ*Moade4* mutant (5×10^4^ spores/mL). Rice leaves sprayed with water (mock) was used as control.

To measure the expression of conidiation-related genes and other genes of *de novo* purine biosynthetic pathway, the RNA samples were from Guy11 and the Δ*Moade4* mutant cultured in liquid CM as described above

Total RNA for all the samples was extracted using Magen universal RNA kit as described previously (Lin et al., 2019). RNAs samples were then reverse transcribed using SYBR® Premix Ex. Taq™ (Tli RNaseH Plus) purchase (Takara Biomedical Technology, Beijing Co. Ltd). qRT-PCR data was generated with Eppendorf Realplex2 mastercycler (Eppendorf AG 223341, Hamburg). Data analysis was by use of delta delta-CT (2−ΔΔCT) method as described by (Livak & Schmittgen, 2001). The expression level of actin was used as the positive control.

### Microscopy

An Olympus DP80 light microscope (Japan) was used to examine conidiophore development, conidia shapes, appressorium formation on inductive surfaces, appressorium penetration, and invasive hyphae development. To observe the fluorescence of MoAde4-GFP, a confocal microscope equipped with Nikon A1 plus instrument (Nikon, Tokyo, Japan) was used.

### Glycogens staining and cytorrhysis assays

The appressorium turgor was measured using an incipient cytorrhysis assay. 20 µl of the conidial suspension (5×10^4^ spores/mL) were placed on hydrophobic coverslips and incubated in a humid chamber for 24 h at room temperature. The water surrounding the conidia was removed carefully and then replaced with an equal volume 20 µl of glycerol in concentrations ranging from 1.0 to 3.0. The number of appressoria that had collapsed after 10 min was recorded. The experiments were repeated three times, and >100 appressoria were observed for each replicates For glycogens staining assay, samples were stained with a solution consisting of 60 mg/ml KI and 10 mg/ml I2 as previously described (Thines et al., 2000).

### Bioinformatic analysis

BLASTp program using *S. cerevisiae* Ade4 amino acid sequence was used to search for MoAde4 in the *M. oryzae* genome database (http://www.kegg.jp/kegg-bin/show_organism?org=mgr). Other fungi ade4 orthologs, were obtained from the National Centre for Biotechnology Information database (NCBI) using MoAde4 as a query. Domains were predicted by Pfam (http://pfam.janelia.org/) while alignment and phylogenetic analysis of the obtained amino acid sequences were performed using MEGA version 6. A phylogenetic tree was generated using the Maximum-Likelihood method, with branches of the tree tested with 1000 bootstrap replicates.

---

**Fig. S1.**
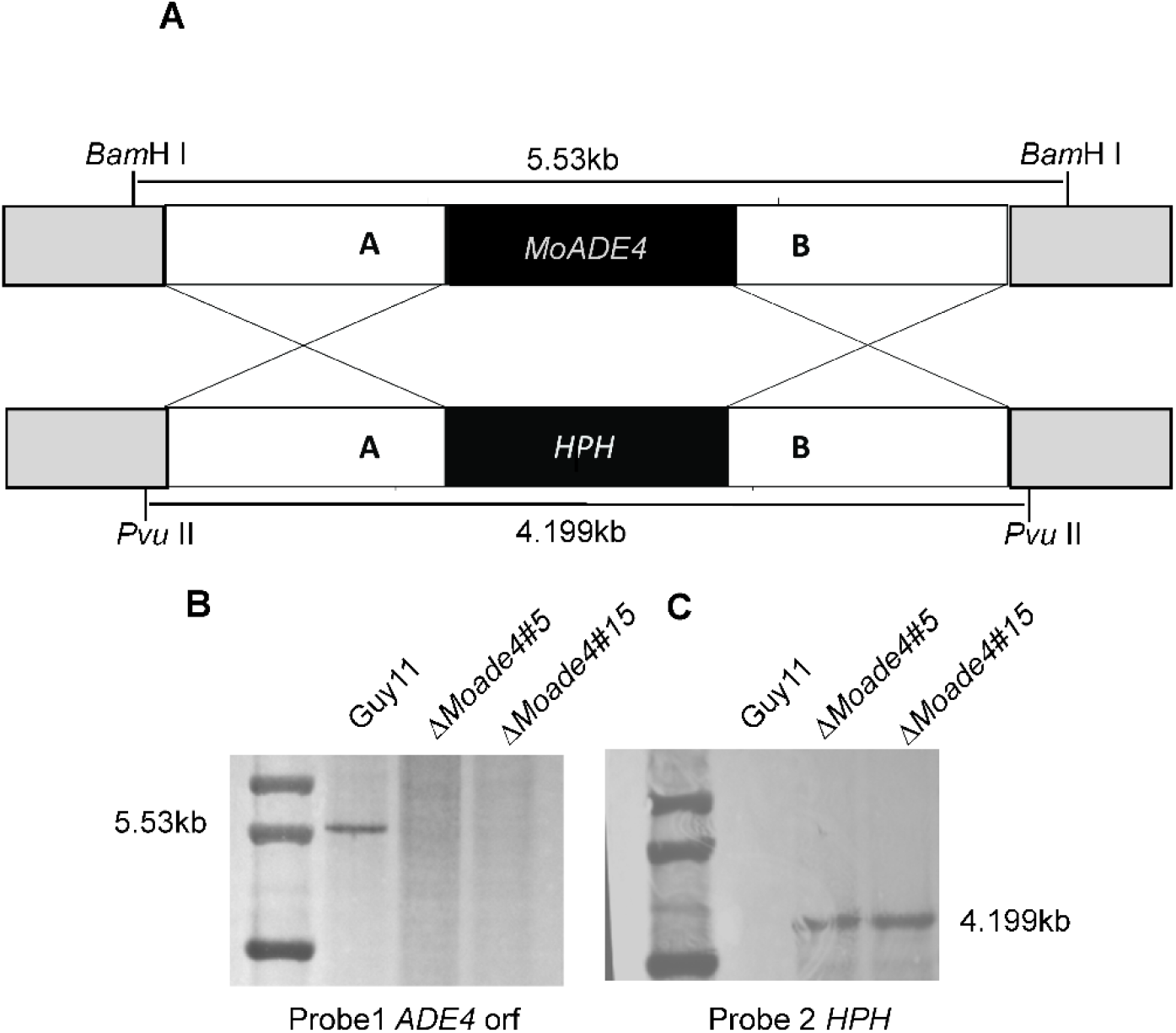
Southern blot analysis of the *MoADE4* gene deletion mutant. (**A**) The strategy of knocking out *MoADE4* gene in the *M. oryzae* genome. (**B**) Southern blot analysis of the WT Guy11 and the gene knockout mutants using *MoADE4* gene (probe 1) (**C**) Southern blot analysis of the gene knockout mutants and WT Guy11 using a hygromycin phosphotransferase (hph) (probe hph).

**Fig. S2.**
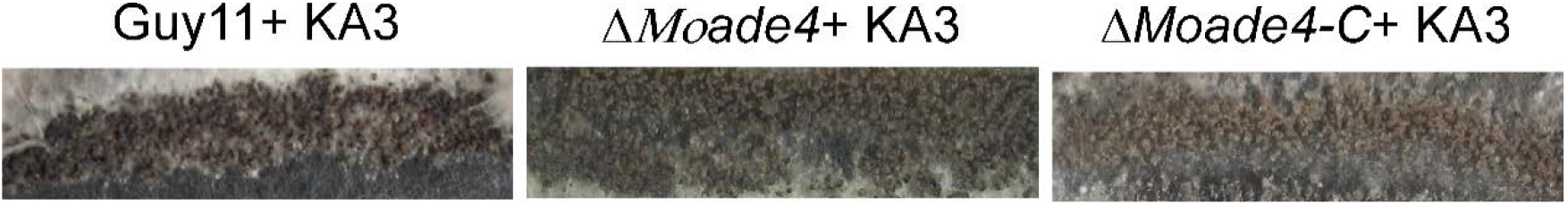
Perithecium production of the wild type Guy11, Δ*Moade4* and the complemented strain on oatmeal agar medium (OTM) after being crossed with KA3. Perithecia formation occurred at the cross point of these fertile strains after 3 weeks of inoculation.

**Table S1.**
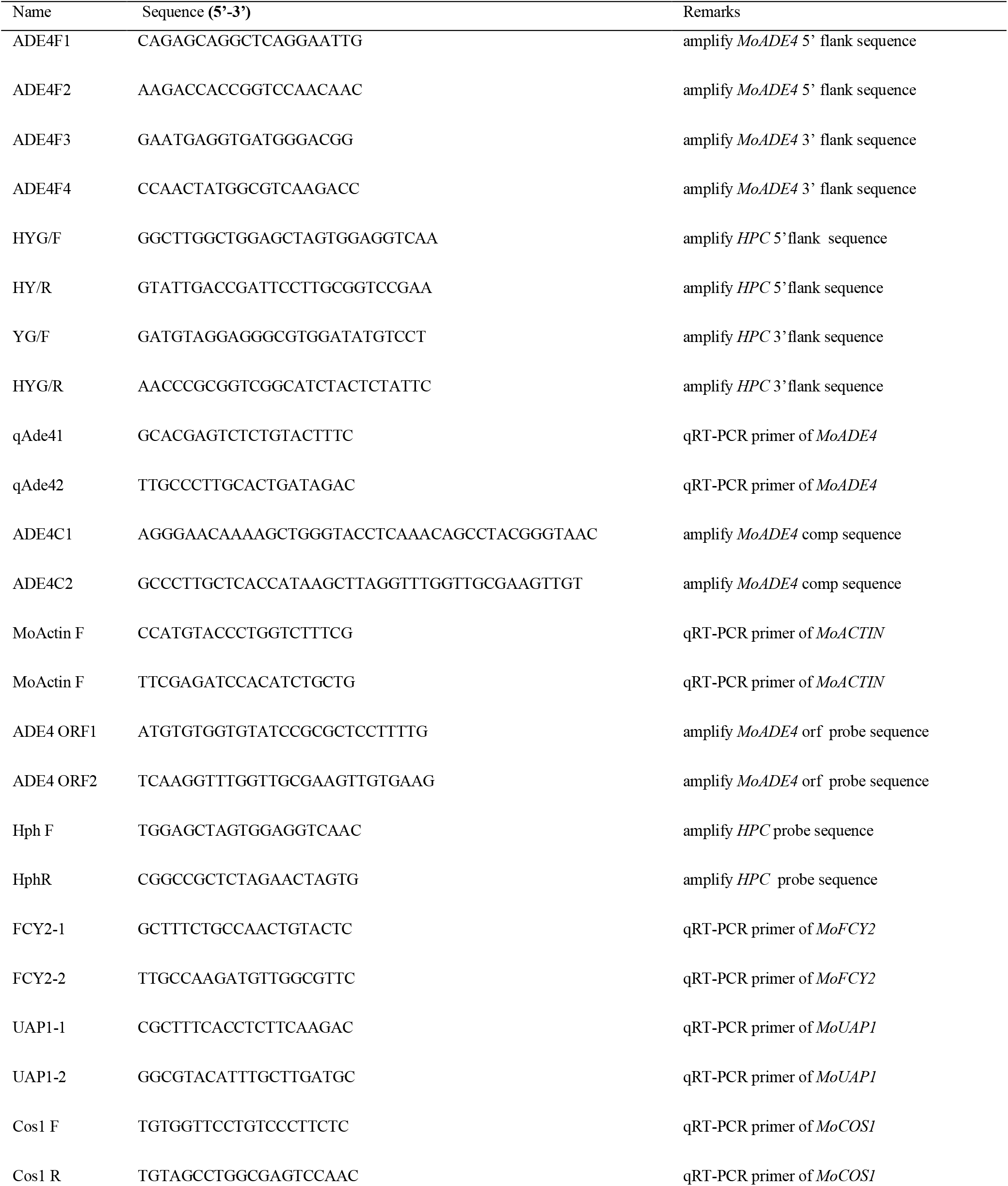
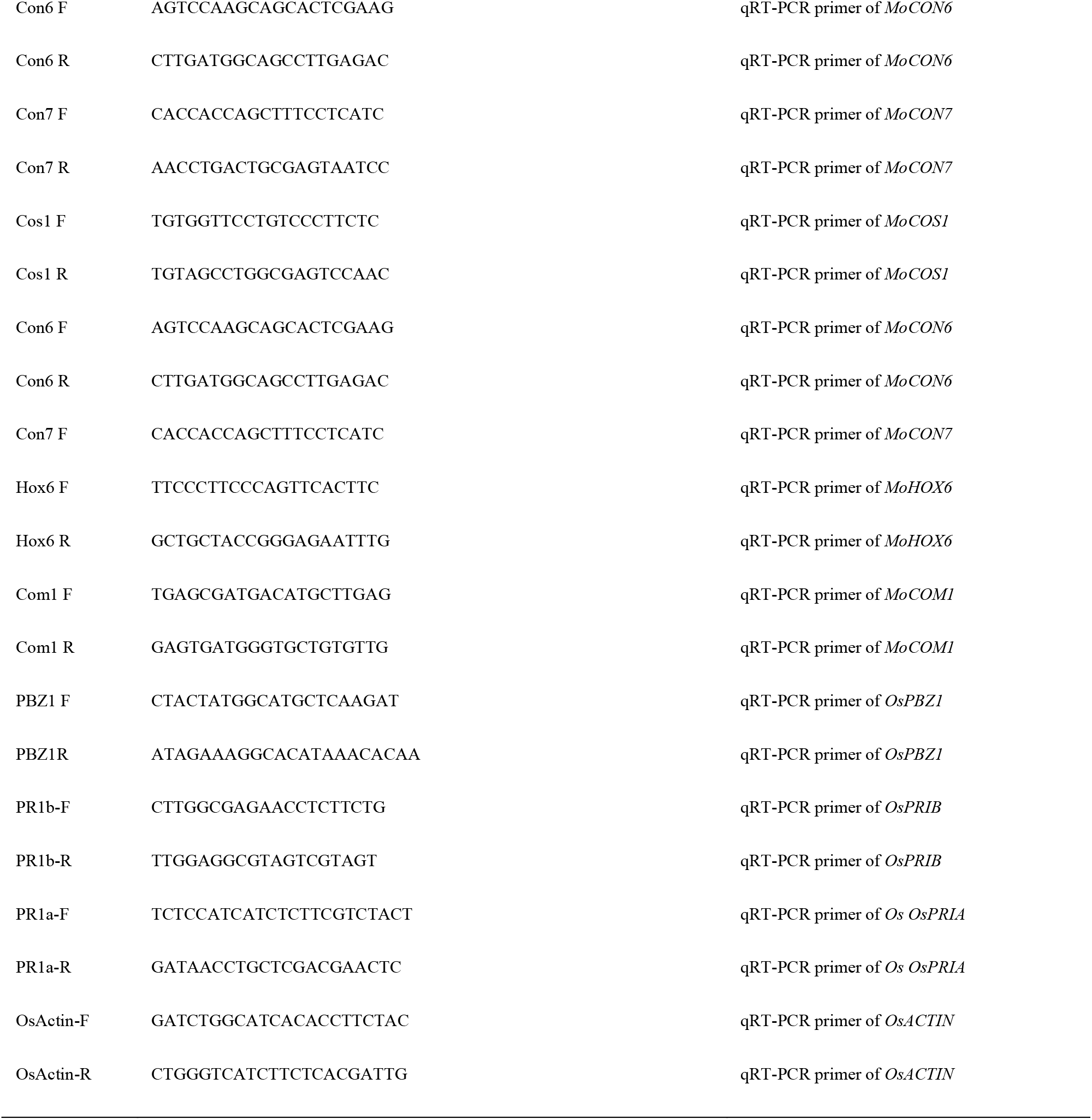
Primers used in this study.

## Acknowledgment

The research was supported by funds from the Natural Science Foundation of China (31601584) and the Science Fund from Distinguished Young Scholars of Fujian Agriculture and Forestry University to W. T. (KXJQ17020). We are grateful to Dr. Anjago Wilfred Mabeche for the advice on the project.

